# A mobile ELF4 delivers circadian temperature information from shoots to roots

**DOI:** 10.1101/612671

**Authors:** Wei Wei Chen, Nozomu Takahashi, Yoshito Hirata, Dmitri A. Nusinow, Steve A. Kay, Paloma Mas

## Abstract

The circadian clock is synchronized by environmental cues, mostly by light and temperature. Elucidating how the plant circadian clock responds to temperature oscillations is crucial to understand plant responsiveness to the environment. Here we found a prevalent temperature-dependent function of the Arabidopsis clock component ELF4 (EARLY FLOWERING 4) in the root clock. Micrografting assays and mathematical analyses show that ELF4 moves from shoots to regulate rhythms in roots. ELF4 movement does not convey photoperiodic information but trafficking is essential to control the period of the root clock in a temperature-dependent manner. At low temperatures, ELF4 mobility is favored and results in a slow-paced root clock while high temperatures decrease movement, leading to a fast clock. Hence, the mobile ELF4 delivers temperature information and establishes a shoot-to-root dialogue that sets the pace of the clock in roots.

## Introduction

Nearly all photosensitive organisms have evolved timing mechanisms or circadian clocks able to synchronize metabolism, physiology and development in anticipation to the 24-hour light/dark cycles^1^. In *Arabidopsis thaliana*, the molecular clockwork is based on complex regulatory networks of core clock components that generate rhythms in a myriad of biological outputs^2, 3^. Appropriate phasing of biological processes relies on clock resetting by light and temperature cues; a mechanism that requires effective changes in the expression and activity of essential clock components^4^. Circadian clocks are also defined by a conserved feature known as temperature compensation^5^. In contrast to the temperature dependency of many physicochemical and biological activities, the circadian clock is able to maintain a constant period over a range of physiological temperatures. By virtue of different transcriptional, post-transcriptional and post-translational mechanisms^6-9^, the plant circadian system buffers the circadian period length. Therefore, the circadian clock does not run faster at higher than at lower temperatures, sustaining a period close to 24-hours within a physiological range of temperatures. An ample collection of light-related factors^10-14^ and clock-associated components^9, 15, 16^ has been shown to directly or indirectly regulate clock temperature compensation in plants.

Among the Arabidopsis clock components, ELF4 (EARLY FLOWERING 4) was initially identified by its role in photoperiod perception and circadian regulation^17^. ELF4 protein assembles into a tripartite complex (Evening Complex, EC) together with ELF3 and LUX ARRHYTHMO or PHYTOCLOCK1 (LUX/PCL1)^18, 19^ to regulate growth and repress circadian gene expression^20, 21^. ELF4 promotes the nuclear localization of ELF3^18^ while LUX directly binds to the promoters of the target genes and thus facilitates the recruitment of ELF4 and ELF3^19, 22^. Loss-of-function mutants of any of the EC components lead to arrhythmia^17, 23-25^. Through multiple interactions with light, clock and photomorphogenesis related factors^26^, the EC is able to coordinate plant responses to environmental cues including temperature^15, 26-30^.

Regarding the circadian structure and organization within the plant, it is broadly accepted that every plant cell harbors a circadian oscillator. However, circadian communication or coupling among cells and tissues varies at different parts of the plant^31-35^. For instance, while cotyledon cells present circadian autonomy^36^, different degrees of cell-to-cell coupling have been reported in leaves^37, 38^, in the vasculature with neighbor mesophyll cells^39^, in guard cells^40^, in cells at the root tip^41, 42^ and within the shoot apex^43^. Long-distance shoot-to-root photosynthetic signaling is also important for clock entrainment in roots^44^ while light piping down the root^45^ contributes to this entrainment. Micrografting assays and shoot excision^43^ suggest the existence of a long-distance mobile circadian signal from shoots to roots. Here we report that ELF4 moves from shoots to control the pace of the root clock in a temperature-dependent manner.

## Results

### Prevalent function of ELF4 sustaining rhythms in roots

We first approached the investigation of the circadian mobile signal by simultaneously following rhythms in shoots and roots of intact plants^43^. The waveforms of the morning-expressed *CCA1* (*CIRCADIAN CLOCK ASSOCIATED 1*) and *LHY* (*LATE ELONGATED HYPOCOTYL*) promoter activities displayed a long period, slightly reduced amplitude and phase delay in roots (Rt) compared to shoots (Sh) (Fig. 1a-b and Supplementary Fig. 1a). The mRNA rhythmic accumulation assayed by Reverse Transcription-Quantitative Polymerase Chain Reaction (RT-QPCR) followed the same trend (Fig. 1c). Similar patterns were observed for the promoter activity of the evening-expressed clock component TOC1/PRR1 (TIMING OF CAB EXPRESSION1/PSEUDO RESPONSE REGULATOR1) (Supplementary Fig. 1b-c). Therefore, the clock is fully operative in roots but its overall pace is slower and the phase delayed compared to shoots.

**Fig. 1.**
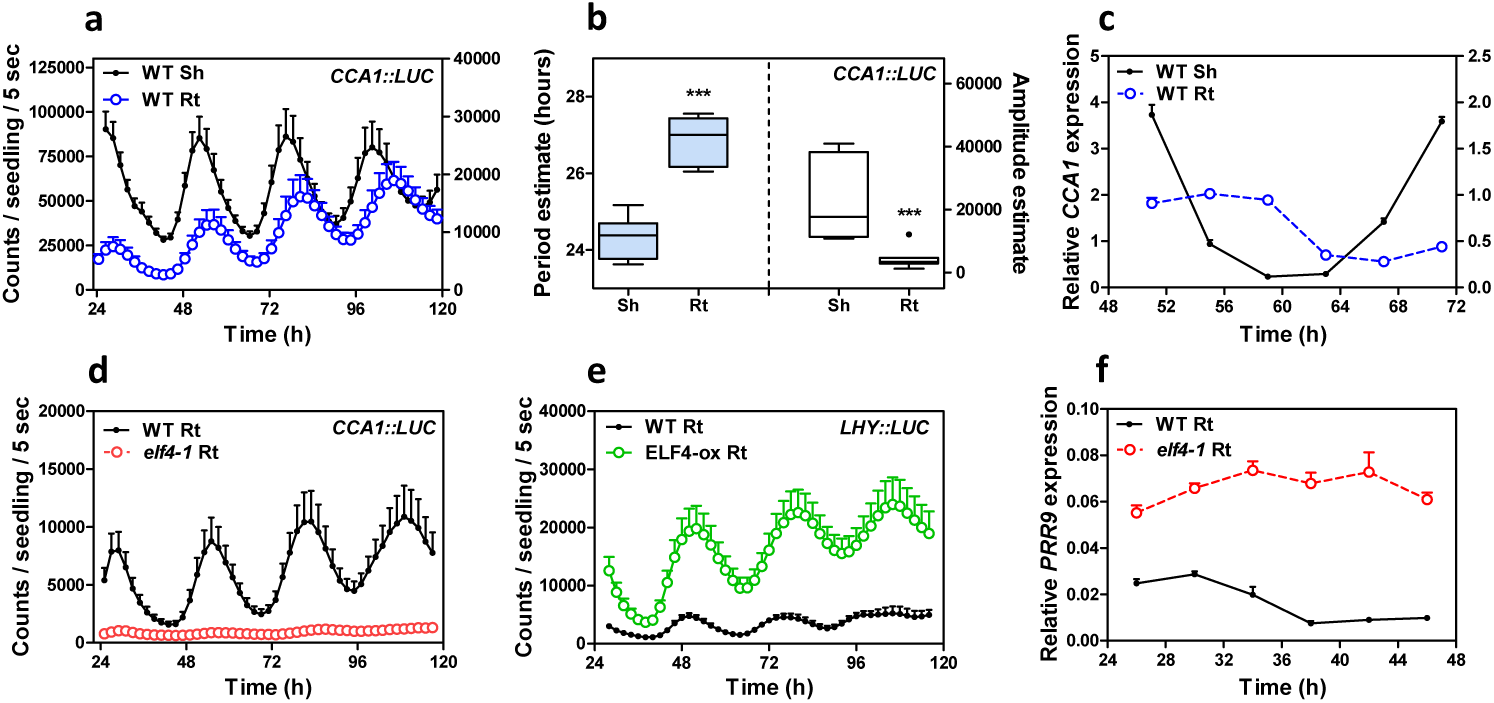
Prevalent function of ELF4 sustaining circadian rhythms in roots. **a,** Luminescence of *CCA1∷LUC* (*LUCIFERASE*) oscillation simultaneously measured in shoots (Sh) and roots (Rt). Root luminescence signals are represented in the right Y-axis. **b**, Period (left Y-axis) and amplitude (right Y-axis) estimates of *CCA1∷LUC* rhythms. *** p-value<0.0001. **c**, Circadian time course analyses of *CCA1* mRNA expression in WT Sh and Rt. **d**, Luminescence of *CCA1∷LUC* rhythms in WT and *elf4-1* Rt. **e**, Luminescence of *LHY∷LUC* rhythms in WT and ELF4-ox Rt. **f**, Circadian time course analyses of *PRR9* mRNA expression in roots of WT and *elf4-1*. Sampling was performed under constant light conditions (LL) following synchronization under light:dark cycles (LD). Data are representative of at least two biological replicates.

The circadian clock is unable to properly run in mutant plants of any of the EC components^17, 23-25^. We therefore examined the role of the EC components in the root clock, and in particular, we focused on ELF4. Circadian time course analyses in WT and *elf4-1* mutant roots showed the fully suppressed *CCA1* and *LHY* promoter activities and mRNA expression in *elf4-1* (Fig. 1d and Supplementary Fig. 1d-f) in a similar way to that described in shoots^21^ (Supplementary Fig. 1g-i). Conversely, over-expression of ELF4 (ELF4-ox) lengthened the period of *LHY∷LUC* (Fig. 1e) indicating that increased ELF4 activity in roots makes the clock to run slow. The expression of *PRR9 (PSEUDO-RESPONSE REGULATOR 9)*, a previously described direct target of the EC in shoots, was clearly up-regulated in *elf4-1* mutant roots (Fig. 1f) suggesting that the EC also represses *PRR9* in roots. Thus, ELF4 plays an important regulatory function in the root clock: mutation abolishes rhythms while over-expression lengthens the circadian period.

### ELF4 moves from shoots to regulate oscillator gene expression in roots

Our previous study showed that a signal from shoots is important for circadian rhythms in roots^43^. Micrografting assays are a powerful tool to identify the nature of mobile signals. The grafting technique *per se* does not alter the rhythms in roots, as grafted WT scions into WT roots show similar rhythms as non-grafted WT plants (Supplementary Fig. 2a-b)^43^. By micrografting different genotypes, we found that grafts of ELF4-ox shoots into *elf4-1* rootstocks [ELF4-ox(Sh)/*elf4-1*(Rt)] (Supplementary Fig. 2c) were particularly efficient in recovering the rhythms in roots (Fig. 2a and Supplementary Fig. 2d-e). The results are noteworthy as *CCA1∷LUC* rhythms are severely disrupted in *elf4-1* mutant roots (Fig. 1d). Restoration of the rhythms was reflecting the circadian function exclusively in roots as water instead of luciferin was applied to shoots (ELF4-ox, Sh, H2O) to exclude the possibility of luminescence signals leaking from shoots into roots of adjacent wells. Rhythms in roots were also recovered when ELF4-ox scion was grafted into *elf4-2* mutant (Supplementary Fig. 2f) rootstocks (Fig. 2b). The influence of shoots as a driving rhythmic force of *elf4-1* rootstocks was mathematically analyzed with recurrence plots obtained by delay coordinates of the grafting time series (Fig. 2c-d). The waveforms of the driving rhythmic force reconstructed from the driven system and their autocorrelation analyses showed a strong periodicity after grafting (Supplementary Fig. 3a-d). In analyses with 10000 randomly shuffled surrogates using as null hypothesis of no serial dependence, we obtained a p-value of 0.2341 (black dash line) (Fig. 2e) before grafting and 0.0004 (gray dash line) after grafting (Fig. 2f). The statistics suggest that after grafting, roots are being forced by a periodic signal from shoots.

**Fig. 2.**
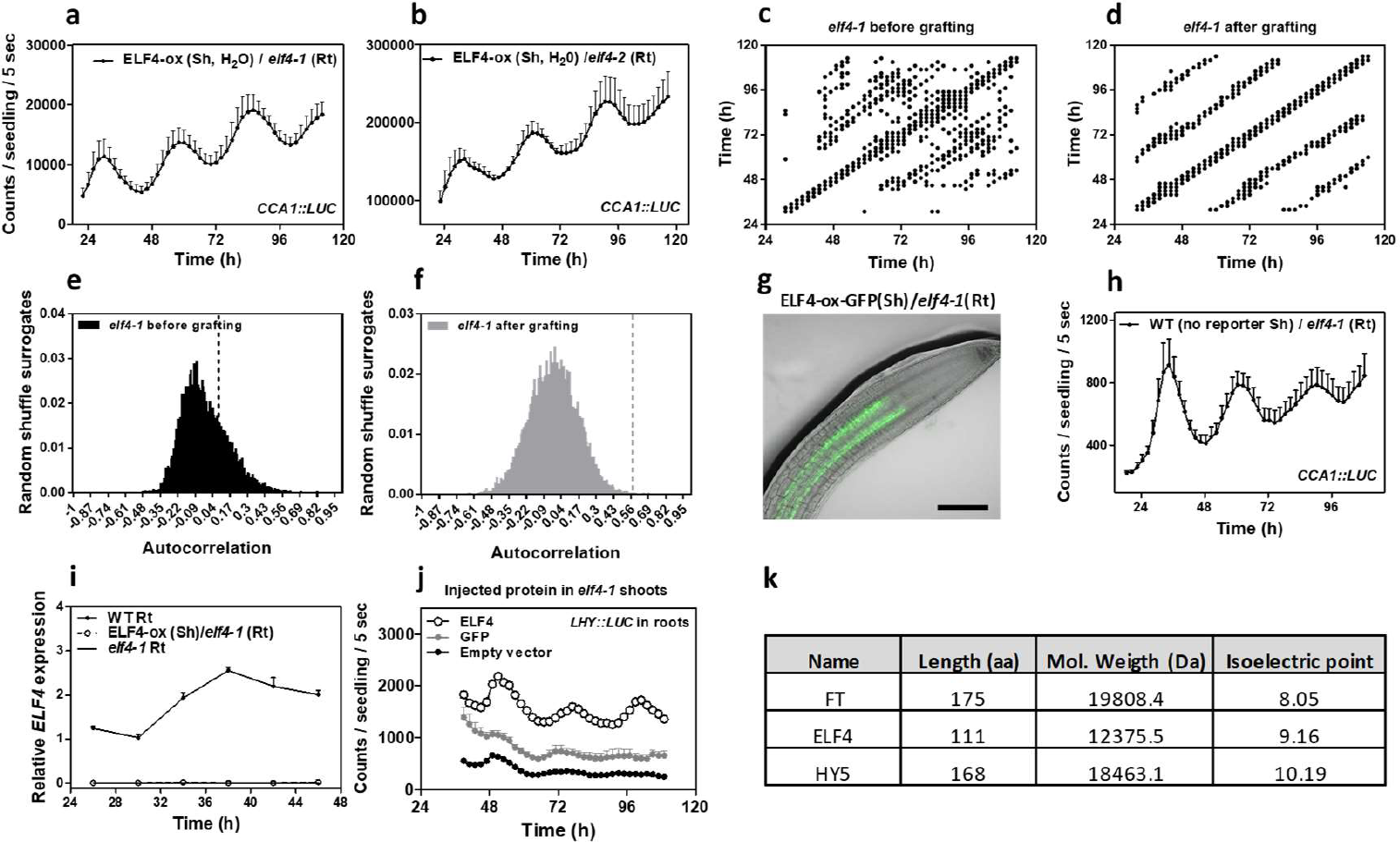
ELF4 moves from shoots to regulate oscillator gene expression in roots. *CCA1∷LUC* luminescence in roots of ELF4-ox scion with **a**, *elf4-1* and **b**, *elf4-2* rootstocks. Water instead of luciferin was added to the wells containing ELF4-ox shoots. **c, d**, Recurrence plots for the driver obtained by delay coordinates and **e,** autocorrelation analyses before **c, e**, and after **d, f**, ELF4-ox scion grafting into *elf4-1* rootstock. **g**, Representative image showing fluorescence signals in roots of ELF4-ox scion and WT rootstock. Scale bar: 100 μm. **h**, *CCA1∷LUC* luminescence in roots of WT scion and *elf4-1* rootstocks. **i**, Circadian time course analyses of *ELF4* mRNA expression in roots of WT, *elf4-1* and ELF4-ox scion and *elf4-1* rootstocks. **j**, Luminescence of *LHY∷LUC* rhythms in *elf4-1* roots after injection of purified ELF4 or GFP proteins in shoots. **k**, Protein features of various plant mobile proteins. At least two biological replicates were performed per experiment.

**Fig. 3.**
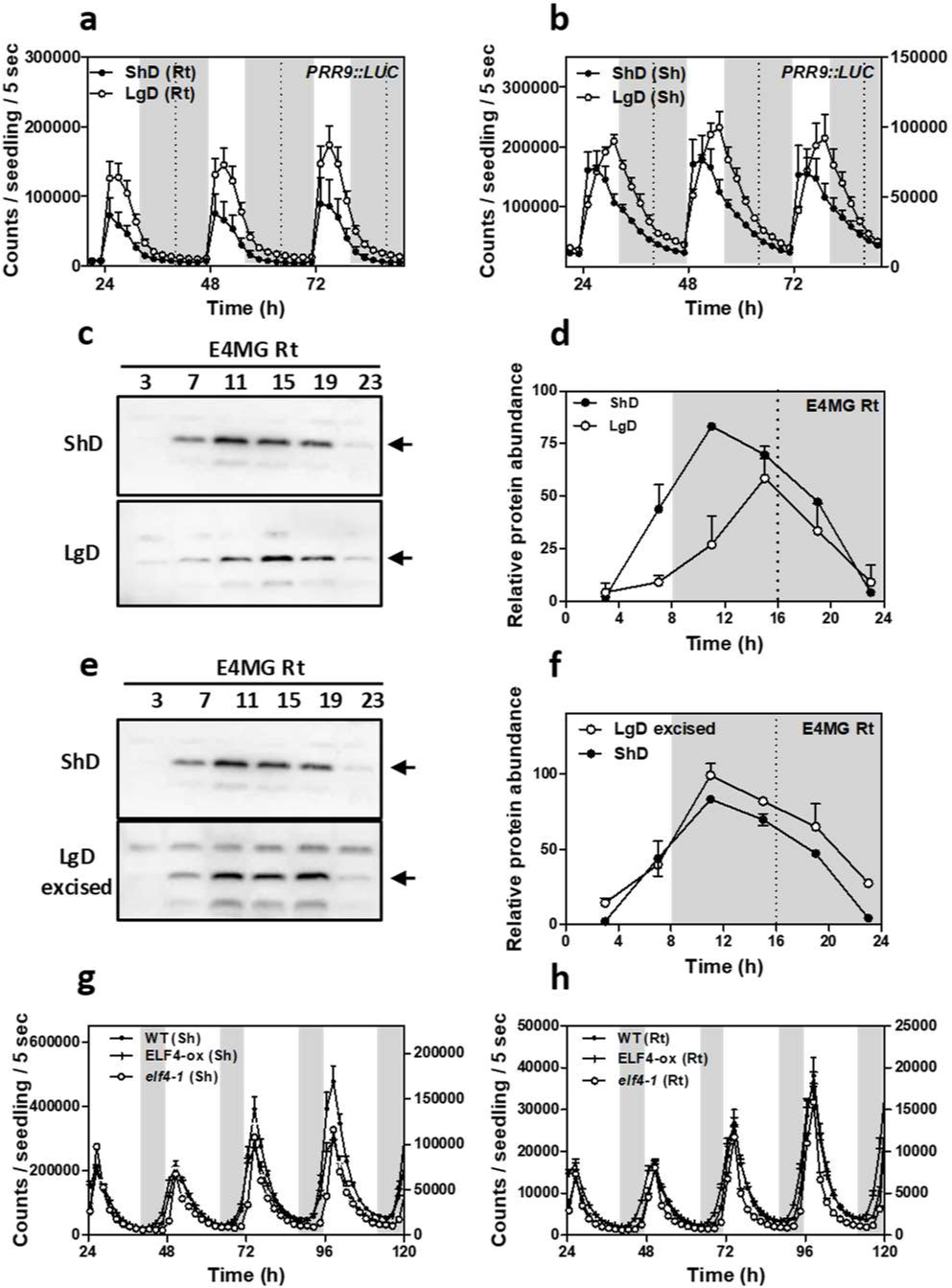
Mobile ELF4 does not regulate the photoperiodic-dependent phase in roots. Luminescence analyses of *PRR9∷LUC* rhythms in **a**, roots and **b**, shoots of plants grown under short day (ShD) or long day (LgD) conditions. **c**, Western-blot analyses and **d**, quantification of ELF4 protein accumulation in ELF4 Minigen roots (E4MG Rt) of plants grown under ShD and LgD (also in Supplementary Fig. 6a-d). **e**, Western-blot analyses and **f**, quantification of ELF4 protein accumulation in E4MG roots of plants grown under ShD and excised roots under LgD (also in Supplementary Fig. 6a-f). Arrows indicate the ELF4 protein. Luminescence of *LHY∷LUC* oscillation in WT, ELF4-ox and *elf4-1* plants measured in **g**, shoots (Sh) and **h**, roots (Rt) under LgD conditions. Dashed lines indicate dusk under LgD. At least two biological replicates were performed per experiment.

Shoots of plants over-expressing ELF4 fused to GFP (GREEN FLUORESCENT PROTEIN) were also grafted into *elf4-1* mutant roots. Confocal imaging showed that ELF4-GFP fluorescent signals accumulated in the vasculature of *elf4-1* mutant rootstock, across the graft junctions (Fig. 2g and Supplementary Fig. 4a-c). To exclude the possibility that the observed results were due to the high over-expression of ELF4-ox plants, we grafted WT shoots into *elf4-1* roots. Although the recovery of the rhythms was not as robust as with ELF4-ox grafts, a rhythmic pattern was observed in roots (Fig. 2h). Thus, ELF4 mRNA or protein are able to move from shoots to roots. This notion was reinforced by the results showing the rhythmic recovery of *elf4-1* rootstocks grafted with ELF4 Minigen (E4MG) scion (Supplementary Fig. 2g-i). To investigate whether the mRNA could be the mobile signal, we performed RT-QPCR time course analyses of roots from ELF4-ox (Sh)/*elf4-1*(Rt) grafts. Our results showed no detectable amplification of *ELF4* mRNA at any time point analyzed (Fig. 2i), which suggest that *ELF4* mRNA might not move through the graft junctions. To confirm this notion, we injected purified ELF4 protein into *elf4-1* mutant (Supplementary Fig. 4d-f). Injection in shoots was able to restore rhythms in roots (Fig. 2j). Although the percentage of ELF4-injected plants with recovered rhythms was low (about 5-8%), the fact that rhythms were actually restored is supportive of a mobile ELF4 protein from shoots to roots. Rhythmic recovery was not apparent when purified GFP was injected (Fig. 2j). Other mobile proteins such as FT (FLOWERING LOCUS T), and HY5 (LONG HYPOCOTYL 5) share some features with ELF4 protein in terms of low molecular weight and high isoelectric point (Fig. 2k).

**Fig. 4.**
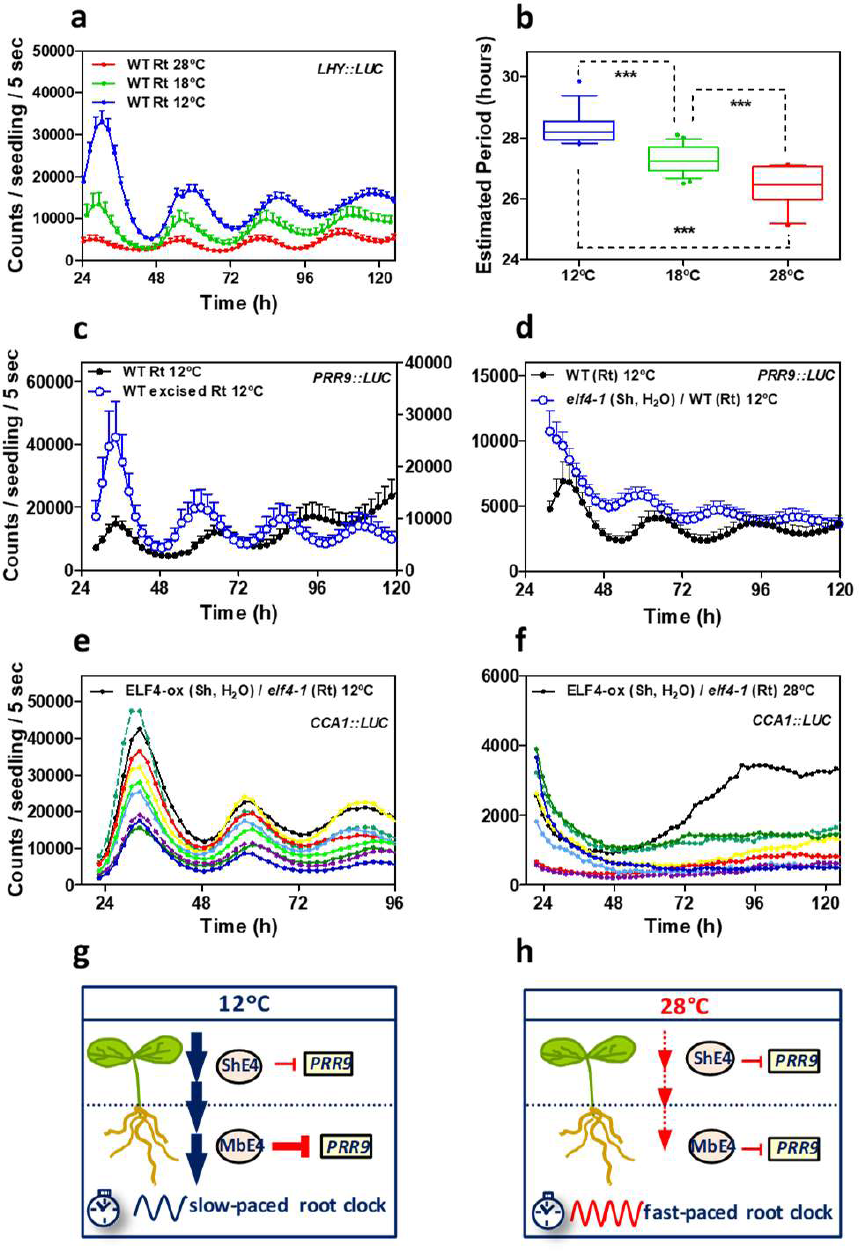
Mobile ELF4 sets the temperature-dependent pace of the root clock. **a**, Luminescence waveforms and **b**, circadian period estimates of *LHY∷LUC* rhythmic oscillation in WT roots at 28°C, 18°C and 12°C. **c**, Luminescence waveforms of *PRR9∷LUC* rhythmic oscillation in WT intact and WT excised roots at 12°C. **d**, *PRR9∷LUC* luminescence in roots of *elf4-1* scion into WT rootstocks at 12°C. Water instead of luciferin was added to the wells containing *elf4-1* scions. Circadian period estimates of ELF4-ox roots at 12°C, 18°C, 28°C and excised roots at 28°C. **e**, Individual luminescence waveforms of *CCA1∷LUC* rhythmic oscillation in ELF4-ox scion into *elf4-1* rootstocks at **e**, 12°C and **f,** 28°C. Water instead of luciferin was added to the wells containing ELF4-ox scions. Schematic drawing depicting the increased ELF4 movement and slow pace of the root clock **g**, at low temperatures and **h**, the decreased movement and fast-paced root clock at high temperature. *** p-value<0.0001; At least two biological replicates were performed per experiment.

### Blocking ELF4 movement by shoot excision alters circadian rhythms in roots

We next attempted to unveil the effects of the mobile ELF4 by blocking ELF4 movement through shoot excision. Analyses of the rhythms showed that excised roots sustained robust oscillations (Supplementary Fig. 5a-b) confirming that the root clock is able to run in the absence of shoots. However, comparison of intact versus excised roots uncovered a shorter period and advanced phase in excised roots (Supplementary Fig. 5c-d). As accumulation of ELF4 results in long periods in shoots^21^ and roots (Fig. 1e), it is plausible that blocking ELF4 movement by shoot excision leads to shorter periods in excised roots. If that is the case, blocking ELF4 movement should also affect ELF4 target gene expression in excised roots. Time course analyses by RT-QPCR revealed that the expression of *PRR9* and *PRR7* was up-regulated in excised roots compared to intact roots (Supplementary Fig. 5e-f), which suggest that in the absence of ELF4 movement from shoots, repression of these genes is alleviated in roots. The use of ELF4-ox intact roots confirmed that *PRR9* and *PRR7* are targets of ELF4 as their expression was clearly down-regulated in intact ELF4-ox roots compared to WT intact roots (Supplementary Fig. 5g-h).

### Mobile ELF4 does not regulate the photoperiodic-dependent phase in roots

In aerial tissues, the circadian clock controls the photoperiodic regulation of growth and development^46^. To determine whether ELF4 movement is important to deliver photoperiodic information, we analyzed rhythms under short day (ShD) and long day (LgD) conditions. In roots, *PRR9∷LUC* waveforms displayed a subtle phase delay under LgD compared to ShD (Fig. 3a) following a similar trend to that observed in shoots (Fig. 3b). Time course analyses by Western-blot of roots of ELF4 Minigen plants^20^ confirmed the phase delay of ELF4 protein accumulation under LgD compared to ShD (Fig. 3c-d). We reasoned that if ELF4 movement is correlated with the photoperiodic-dependent phase delay, then excision of shoots might affect the phase shift in roots. In agreement with the oscillations in promoter activity and gene expression (Supplementary Fig. 5c-d), the phase of ELF4 protein accumulation was advanced following excision under both LgD and ShD (Supplementary Fig. 6a-d). Interestingly, under LgD conditions, excision rendered a similar pattern of ELF4 accumulation than in intact roots under ShD (Fig. 3e-f). Therefore, excision abolished the phase delay observed in intact root under LgD (compare Fig. 3c-d with Fig. 3e-f). The results suggest that the photoperiodic-dependent phase shift in roots is hampered by blocking ELF4 movement. However, excised roots still showed the phase delay under LgD compared to excised roots under ShD (Supplementary Fig. 6e-f). Furthermore, analyses of rhythms under LgD conditions showed that plants miss-expressing ELF4 (ELF4-ox and *elf4-1* mutant) displayed very similar rhythms to WT both in shoots and roots (Fig. 3g-h) suggesting that ELF4 function is not essential to sustain rhythms under entraining conditions. Together, the results suggest that blocking ELF4 movement by excision advances the phase of the root clock but the mobile ELF4 does not directly regulate the photoperiodic-dependent phase shift in roots.

### ELF4 movement contributes to the temperature-dependent changes in circadian period of the root clock

As the EC also coordinates temperature responses, we examined whether a mobile ELF4 can convey temperature information from shoots to roots. To that end, we first examined the effect of different temperatures (28°C, 18°C and 12°C) on circadian rhythms in roots. We found that *LHY∷LUC* circadian period length was significantly shorter at high than at low temperatures (Fig. 4a-b). Shortening of period length at increasing temperature was also observed for other circadian reporter lines (Supplementary Fig. 7a-d). Thus, under our conditions, the circadian clock in roots is not able to sustain circadian period length within a range of temperatures. Temperature compensation, one of the defining properties of circadian systems, does not function in the root clock.

As ELF4 accumulation lengthens period length, we next examined the possible contribution of ELF4 to the long period phenotype at low temperatures. Changes in period length could be mediated by increased ELF4 activity and/or by the increased protein movement from shoots to roots. To examine these possibilities, we compared the effects of blocking ELF4 movement by excision and by grafting at low and high temperatures. Essentially, if the long period in roots at 12°C is independent of movement but results from the increased activity of ELF4, blocking movement from shoots by excision or grafting should not have a major effect on period length. However, if ELF4 movement contributes to the period regulation, abolishing ELF4 traffic should lead to an observable and differential effect on period length at different temperatures.

Our results showed that excision shortened the period length in WT roots and this effect was highly significant at 12°C as compared to the minor effect at 28°C (Fig. 4c and Supplementary Fig. 8a-c). Grafting *elf4-1* scion into WT rootstock resulted in a similar pattern to that of excised WT roots at 12°C (Fig. 4c and d). Therefore, blocking ELF4 movement by excision or grafting shortens the long period of WT roots at 12°C. Analyses of other circadian reporter lines and at 18°C also showed that excision significantly shortened period length compared to intact roots (Supplementary Fig. 8d-e). The results suggest a prevalent function of ELF4 movement in the control of period length at lower temperatures. Interestingly, *elf4-1* grafting into WT rootstock did not noticeably affect the rhythms of WT roots at 28°C (Supplementary Fig. 8f) consistent with the results of WT excised roots at 28°C (Supplementary Fig. 8a-b). The results suggest that ELF4 movement is controlled by temperature. To further verify this notion, we examined rhythmic recovery in grafts of ELF4-ox scion into *elf4-1* rootstock at low and high temperatures and found evident rhythms at 12°C but not at 28°C (Fig. 4e-f). Therefore, ELF4 movement contributes to the temperature-dependent control of circadian period in roots.

Altogether, we propose a model by which mobile ELF4 (mbE4) from shoots to roots defines a pool of active ELF4 protein that is competent to repress target circadian gene expression in roots. ELF4 trafficking is favored at low temperatures, which results in a slow-paced clock (Fig. 4g) while high temperatures decrease the movement, leading to a fast root clock (Fig. 4h). The temperature-dependent movement of ELF4 allows a shoot-to-root dialogue that controls the pace of the clock and provides a mechanism by which temperature cues from shoots set the circadian period length in roots.

## Discussion

The simultaneous examination of rhythms in shoots and roots of single individual plants shows that the promoter activities and mRNA accumulation of clock genes in roots display a longer period and delayed phase compared to shoots. The trend was observed for morning- and evening-expressed key oscillator genes suggesting that the overall circadian system in roots is not as precise as in other parts of the plant (e.g. the shoot apex)^43^. Despite the long period and delayed phase, the rhythms persist in roots for several days under LL, which is reminiscent of a fully functional clock. The lack of precision might provide circadian flexibility for rapid adjustments and improved responses in roots.

The EC directly represses *PRR9* and *PRR7* expression^18, 22, 28, 47, 48^ and indirectly promotes the expression of the morning-expressed oscillator genes *CCA1* and *LHY* ^47-50^. Our analyses with *elf4-1* mutant and ELF4-ox plants demonstrate that ELF4 function in roots is also important for proper repression of *PRR9* and *PRR7* and activation of *CCA1* and *LHY*. ELF4 regulatory function in roots is thus similar to that previously described for the EC using whole plants. Over-expression of ELF4 further delays the phase and lengthens the period of the root clock suggesting that ELF4 slows down the circadian period in roots as in shoots^21^. The fact that accumulation of ELF4 lengthens the period agrees with the results showing that blocking movement by shoot excision or grafting shortens the period while the reduced ELF4 movement at high temperature also leads to shorter periods in roots.

Micrografts of ELF4-ox scion into *elf4-1* or *elf4-2* rootstocks allow a remarkable recovery of rhythms and the accumulation of ELF4-GFP fluorescent signals in the vasculature of *elf4-1* mutant rootstock. These results suggest that ELF4 moves from shoots to roots, which is in agreement with the assays of ELF4 protein injection in shoots and the subsequent rhythmic recovery in roots. The effects are not due to the over-accumulation of ELF4-ox as micrografts of E4MG and WT plants are also able to recover the rhythms of the *elf4-1* mutant roots. Furthermore, our experiments adding water to the scion exclude the possibility that rhythms in grafted roots are due to leakage for the adjacent well containing the shoot. The fact that ELF4 protein shows similar properties in terms of length, molecular weight and isoelectric point to other mobile proteins^52-55^ also support our conclusions of an ELF4 movement.

Excision blocks ELF4 movement from shoots and consequently, we observe that oscillator gene expression is affected in WT excised roots. The phase shifts observed following excision prompted us to examine whether ELF4 movement contributed to the photoperiodic-dependent phase shift. However, excised roots still sustained the phase delay under LgD suggesting that other factors are responsible for this regulation. Light piping down the root^45^ might be important for this synchronization. Regardless the mechanism, it is able to overcome the miss-expression of ELF4 in shoots and roots as ELF4-ox and *elf4-1* mutant plants displayed similar rhythms to WT. Clear alteration of circadian expression under LL but not under entraining conditions has been reported for other clock mutants and over-expressing plants^56^.

As photoperiod does not control ELF4 movement, we focused on temperature. The EC activity is down-regulated at high temperatures in whole seedlings^28, 30^. Shoot excision and micrografting *elf4-1* scion into WT rootstock shortened the period indicating that ELF4 movement is important in the control of circadian period length. Period shortening is more significant at low than at high temperatures confirming that ELF4 movement is favored at low temperatures. As ELF4 accumulation results in long period, the increased movement leads to a clock that runs slower at low than at high temperatures. Therefore, a mobile ELF4 contributes to the lack of temperature compensation in roots. It would be interesting to elucidate whether period sensitivity to temperature might provide an advantage for optimal root responsiveness to temperature variations.

## Materials and Methods

### Plant material and growth conditions

*Arabidopsis thaliana* seedlings were stratified at 4°C in the dark for 2-3 days on Murashige and Skoog (MS) agar medium without or with 3% of sucrose (MS or MS3, respectively) as specified for each experiment. Plates were transferred to chambers with light-and temperature-controlled conditions with 25-40 μmol·quanta·m^−2^·s^−1^ of cool white fluorescent light. Seedlings were synchronized under Light:Dark cycles, LD (12h light: 12h dark) at 22°C. For experiments with different temperatures, seedlings were analyzed under constant light conditions at 12°C, 18°C, 22°C or 28°C following synchronization under LD (12h light: 12h dark) at 22°C. For experiments with different photoperiods, seedlings were grown under short days (ShD, 8h light: 16h dark) or long days (LgD, 16h light: 8h dark). Reporter lines *CCA1∷LUC*^57^, *LHY∷LUC*^18^, *PRR9∷LUC*^58^, *TOC1∷LUC* ^6^ and *elf4-1*^17^, *elf4-2* ^26^, ELF4 Minigene ^19^ and ELF4-ox and ELF4-GFP-ox ^18, 19^ plants were described elsewhere. Plants were transformed using *Agrobacterium tumefaciens* (GV2260)-mediated DNA transfer ^61^. For in vitro protein injection assays, the ELF4 CDS was subcloned into the pET MBP_1a vector (Novagen) after removing the Green Fluorescent Protein CDS, by Nco I and Xho I restriction enzyme digestion.

### In vivo luminescence assays

In vivo luminescence assays were performed as previously described ^43^. Briefly, 7-15 day-old seedlings synchronized under LD cycles at 22°C were transferred to 96-well plates and released into the different conditions as specific for each experiment. Analyses were performed with a LB960 luminometer (Berthold Technologies) using the Microwin software (Mikrotek Laborsysteme). The period, phase and amplitude were estimated using the Fast Fourier Transform–Non-Linear Least Squares (FFT-NLLS) suite ^62^ using the Biological Rhythms Analysis Software System (BRASS, http://www.amillar.org). For the simultaneous analysis of rhythms of shoots and roots from the same plant, the connection between the two adjacent wells of the 96-well plates was serrated. Seedlings were then horizontally positioned so the shoot was placed in one well and the roots in the contiguous well. For excision analyses, roots were excised from shoots and placed into the 96-well plates for luminescence analyses. Data from samples that appeared damaged or contaminated were excluded from the analysis. For analyses of grafted samples, water instead of luciferin was applied to the wells containing shoots to avoid possible leaking signals from shoots to roots as specified. At least two biological replicates were performed per experiment. Each biological replicate included 6 to 12 seedlings per condition and/or genotype.

### Protein purification and injection analyses

*E. coli* cells (BL21, Dh5α) were transformed and grown in LB medium (Tryptone 10 g/L, yeast extract 5 g/L, NaCl 10 g/L pH 7.5) until OD600 values of 0.8-0.10. Isopropyl β-D-1-thiogalactopyranoside (IPTG)-mediated induction of MBP-ELF4 and MBP-GFP was performed at 28°C for 6 h. Bacteria were lysed by sonication for 2-3 minutes (30s on, 30s off, high intensity) using a sonicator (Bioruptor, Diagnode). Recombinant proteins were purified using gravity flow columns with amylose resin (New England Biolabs). MBP cleavage was performed by incubation in cleavage buffer (50 mM Trizma-HCl, pH 8.0, 0.5 mM EDTA, and 1 mM DTT) for 2 hours at 30°C with native Tobacco Etch Virus (TEV) protease (Sigma-Aldrich). The purified recombinant proteins were concentrated using Amicon centrifugal filters following the manufacturer recommendations (Millipore). Protein yield was estimated by measuring absorbance at 595 nm using a spectrophotometer (UV-2600, SHIMADZU). Proteins were also examined by Coomassie-Brilliant Blue staining of polyacrylamide gels to confirm protein size and integrity. Purified ELF4 was injected into leaves of 15-day old *elf4-1* mutant seedlings harboring the *LHY∷LUC* reporter line. Similar concentration of GFP protein was also injected as a negative control. Rhythms were subsequently examined in a LB960 luminometer (Berthold Technologies) as described above.

### Time course analyses of gene expression by RT-qPCR

Seedlings were synchronized under LD cycles in MS3 medium plates for 8-14 days and subsequently transferred to LL. Shoots are roots from intact plants were taken every 4 hour over the circadian cycle. For excised roots, shoots and roots were carefully separated with a sterile razor blade and the excised roots were deposited on MS3 agar medium plates for 2 or 3 days as specified. RNA was purified using a Maxwell RSC Plant RNA kit following the manufacturer’s recommendations (Promega). Single-stranded cDNA was synthesized using iScript Reverse Transcription Supermix for RT-qPCR (Bio-Rad). qPCR analyses were performed with cDNAs diluted 50-fold with nuclease-free water using iTag Universal SYBR Green Supermix (Bio-Rad) or Brilliant III Ultra-Fast SYBR Green qPCR Master Mix (Agilent) with a 96-well CFX96 Touch Real-Time PCR detection system (Bio-Rad). Each sample was run in technical triplicates. The expression of *PP2AA3 (PROTEIN PHOSPHATASE 2A SUBUNIT A3,* AT1G13320) or *MON1* (*MONENSIN SENSITIVITY1*, AT2G28390) was used as a control. Crossing point (Cp) calculation was used for quantification using the Absolute Quantification analysis by the 2^nd^ Derivative Maximun method.

### Time course analyses of protein accumulation by Western-blot assays

Approximately 100 mgs of roots of ELF4 Minigene or ELF4-ox plants grown under the specified light and temperature conditions were sampled at the indicated time points. Samples were rapidly frozen with liquid nitrogen and grounded with stainless steel beads (Millipore) in a tissue lyser (QIAGEN, TissueLyser II). Tissue was subsequently resuspended in Protein Extraction Buffer (PEB) containing 50 mM Tris-HCl pH 7.5, 150 mM NaCl, 0.5% NP40, 1 mM EDTA, and protease inhibitors cocktail (1:100) and PMSF (1:1000). Protein extracts were centrifuged at 4 °C, measured for protein concentration using Bradford reagent (Bio-Rad) and normalized to 2 mg/ ml in 4 x SDS loading buffer (250 mM Tris-HCl, pH 6.8, 8% SDS, 0.08% bromophenol blue, 40% glycerol). Samples were run on a 12% gel and analyzed by immunoblotting, fixed 30 min with 0.4% Glutaraldehyde solution (Sigma-Aldrich) and detected with an anti-HA antibody (Roche) (1:2000 dilution) and a goat anti-rat horse peroxidase conjugated secondary antibody (Sigma-Aldrich) (1:4000 dilution). ELF4 protein from ELF4-ox roots was detected using an anti-GFP antibody (Santa Cruz Biotechnology) (1:500 dilution) and an goat anti-mouse horse peroxidase conjugated secondary antibody (1:500 dilution) (Thermo-Fisher). At least two biological replicates were performed per experiment and/or condition.

### Micrografting assays

Micrografting was performed essentially as previously described ^43^. Data from unsuccessful grafted seedlings that failed to properly join together or grafts that were insufficiently clear to be successful were discarded. Approximately 100-150 grafting events were performed for every combination of grafts. The percentage of successfully micrografted plants was about 30-50 % (possibly higher but only the clearly successful grafted plants were taken into account). From the successfully grafted plants, 30-60 % showed different degrees of recovered rhythms. For in vivo luminescence assays, shoots and roots of grafted plants were simultaneously examined using the protocol described above. For ELF4-ox grafts, water instead of luciferin was added to the wells containing shoots to exclude the possibility that recovery of rhythms in roots were due to leaking signals from shoots. Similarly, grafted WT shoots contained no reporter fused to luciferase.

### Reconstruction of driving forces by recurrence plots

Common driving forces were estimated following a several-step procedure. First, we took time differences of consecutive measurements and remove trends. Second, we used delay coordinates ^74, 75^ of 5 dimensional space to obtain a recurrence plot ^76^ so that 5% points, except for the central diagonal line, have points plotted. Third, we took the union for the recurrence plots to infer the recurrence plot of the common driving force ^77^. Fourth, we applied the assumption of continuity and supplied the points at (i,i+1) and (i+1,i) for each i ^78^. Fifth, we applied the method described in ^77^ to convert the recurrence plot of the common driving force into time series. Lastly, we extracted the component corresponding to the largest eigenvalue. The periodicity of the reconstructed common driving force X(t) was evaluated using the autocorrelation function and 10000 random shuffle surrogates ^79^. Here, the null hypothesis was that there was no serial dependence. The autocorrelation with time difference k is the correlation coefficient between X(t) and X(t+k). Thus, it is close to 1 if X(t) and X(t+k) are similar while it is close to 0 if they are not related to each other. If there is a 24h periodicity in the driving force, the autocorrelation with 24h time difference should be a value close to 1.

### Confocal imaging

For *in vivo* confocal imaging, the roots of WT and ELF4-ox (fused to GFP) grafted shoots into *elf4-1* mutants were placed on microscope slides (Sigma). Fluorescent signals were imaged with an argon laser (transmissivity: 40%; excitation: 515 nm; emission range: 530-630 nm) in a FV-1000 confocal microscope (Olympus, Tokyo, Japan) with a 40x/1.3 oil immersion objective. The image sizes were about 640 x 640 (0.497 μm/pixel) and sampling speed of 4 μs/pixel. The results are representative of at least three biological replicates for grafting and about three-four images per grafts.

## Acknowledgments

We thank members of the Mas laboratory for helpful discussion and suggestions. We also thank Prof. S. Davis (University of York) for the ELF4-ox and *elf4-1* seeds and Prof. T. Nakagawa (Shimane University) and Meiji Seika Kaisha, Ltd. for the Gateway vectors. S.A.K. acknowledges support from the National Institute of Health (grant number GM067837). The Mas laboratory is funded by the FEDER/Spanish Ministry of Economy and Competitiveness and by the Generalitat de Catalunya (AGAUR). P.M. laboratory also acknowledges financial support from the CERCA Program/Generalitat de Catalunya and by the Spanish Ministry of Economy and Competitiveness through the “Severo Ochoa Program for Centers of Excellence in R&D” 2016–2019, grant number SEV-2015-0533.

---

**Supplementary Fig. 1.**
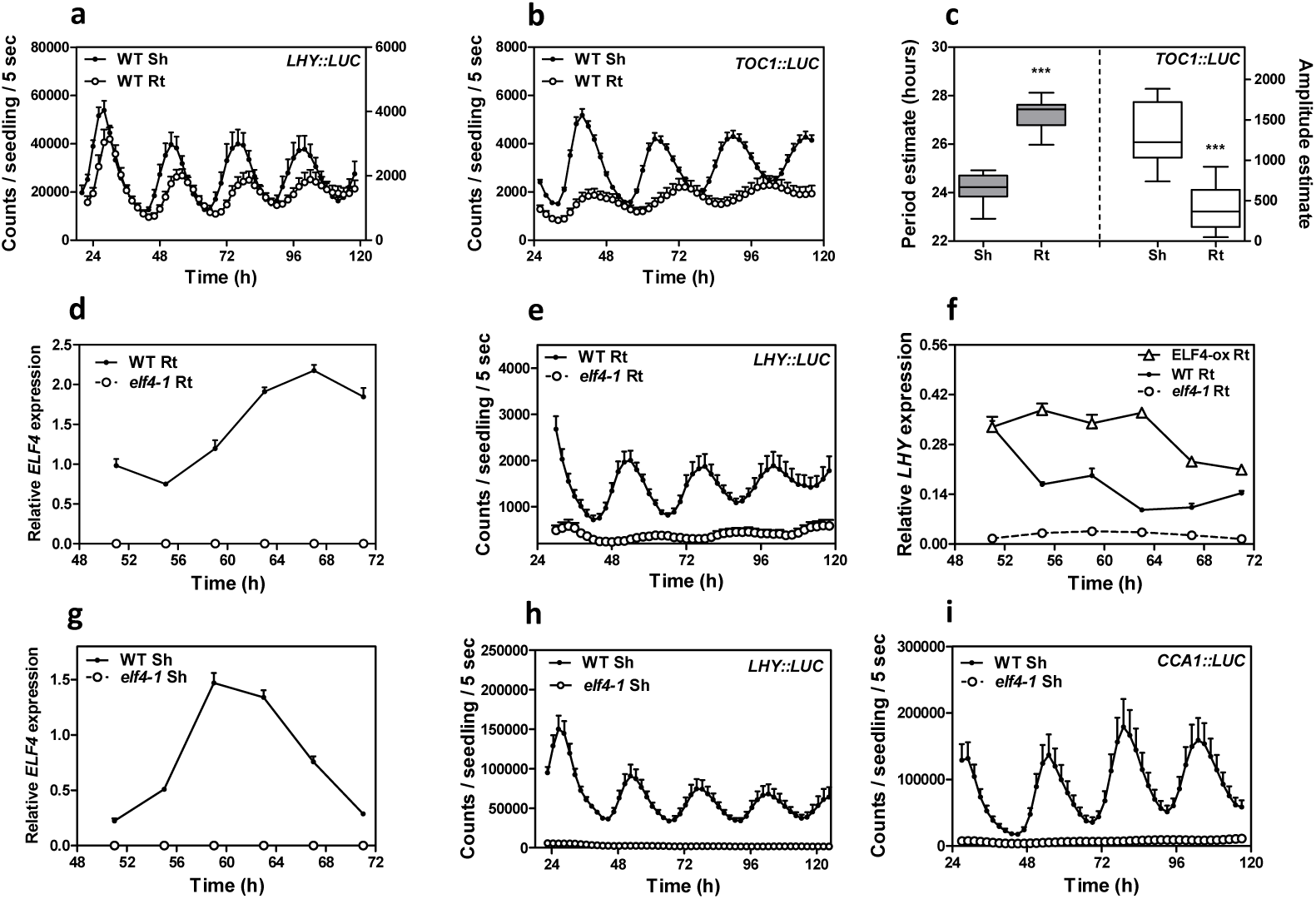
Comparative analyses of rhythmic circadian oscillation in shoots and roots. Luminescence of **a,** *LHY∷LUC*, and **b,** *TOC1∷LU*C rhythms simultaneously measured in shoots (Sh) and roots (Rt). Root luminescence signals in **a,** are represented in the right Y-axis. **c,** Circadian period (left Y-axis) and amplitude (right Y-axis) estimates of *TOC1∷LUC*. **d,** Circadian time course analyses of *ELF4* mRNA expression in WT and *elf4-1* mutant Rt. **e,** Luminescence of *LHY∷LUC* rhythms in WT and *elf4-1* mutant Rt. **f,** Circadian time course analyses of *LHY* mRNA expression in roots of WT, ELF4-ox and *elf4-1*. **g,** Circadian time course analyses of *ELF4* mRNA expression in WT and *elf4-1* mutant Sh (also in Supplementary Fig. 2c). Luminescence of **h,** *LHY∷LUC* and **i,** *CCA1∷LUC* rhythms in WT and *elf4-1* mutant Sh. Each biological replicate included 6 to 12 seedlings per condition and/or genotype. The mRNA expression and promoter activity analyses were performed under constant light conditions previous synchronization of plants under LD cycles at 22°C. At least two biological replicates were performed per experiment.

**Supplementary Fig. 2.**
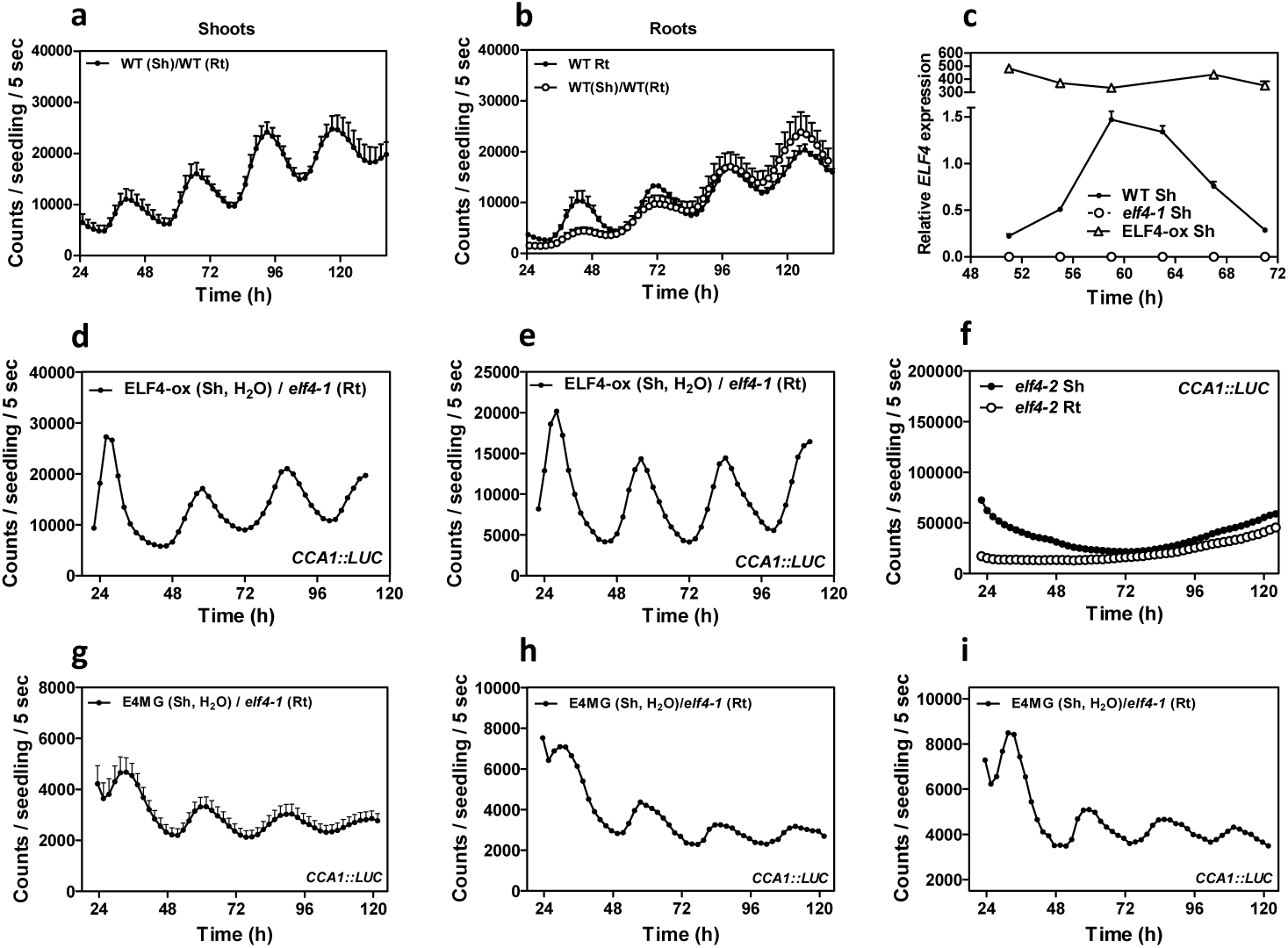
Analyses of circadian rhythmicity of the micrografting assays. *ELF4∷LUC* luminescence in **a,** shoots and **b,** roots of WT scion into WT rootstocks and its comparison with luminescence in (non-grafted) WT roots. **c,** Circadian time course analyses of *ELF4* mRNA expression in shoots of WT, ELF4-ox and *elf4-1* (also in Supplementary Fig. 1g). **d, e,** Individual waveforms of *CCA1∷LUC* rhythmic recovery in roots of ELF4-ox scion and *elf4-1* rootstocks. Water instead of luciferin was added to the wells containing ELF4-ox shoots. **f,** *CCA1∷LUC* luminescence in shoots and roots of *elf4-2* mutant plants. **g,** *CCA1∷LUC* luminescence in roots of ELF4 Minigen (E4MG) scion into *elf4-1* rootstocks. **h, i,** Individual waveforms of *CCA1∷LUC* rhythmic recovery in roots of E4MG scion into *elf4-1* rootstocks. Water instead of luciferin was added to the wells containing ELF4-ox shoots. The mRNA expression and promoter activity analyses were performed under constant light conditions previous synchronization of plants under LD cycles at 22°C. At least two biological replicates were performed per experiment.

**Supplementary Fig. 3.**
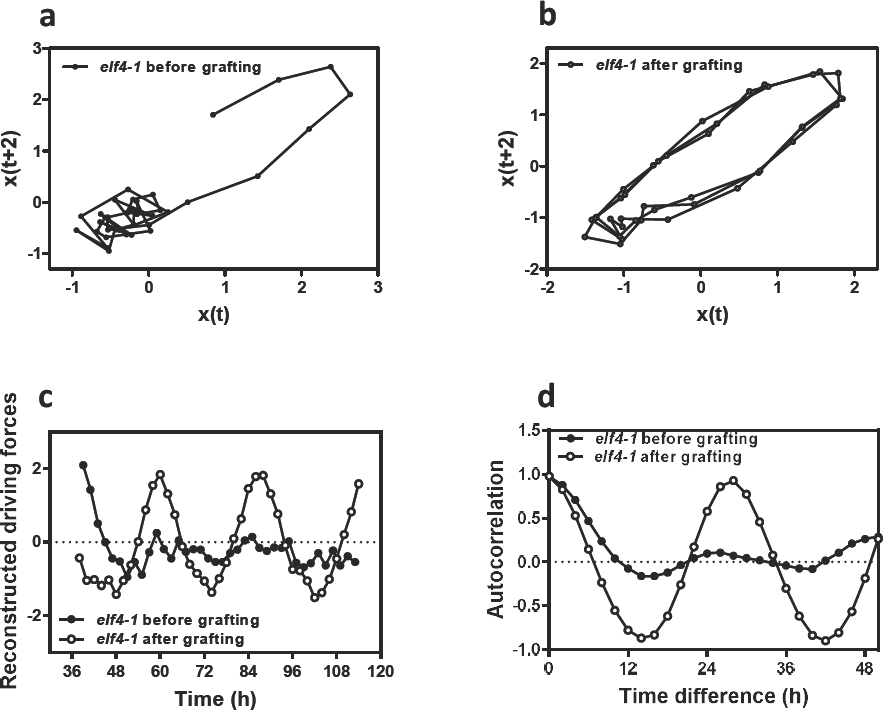
Mathematical analyses of the micrografting effects on rhythms. Two-dimensional plots of *CCA1∷LUC* rhythms in roots before **a,** and after **b,** grating ELF4-ox scion into *elf4-1* rootstocks. **c,** Reconstructed waveforms and **d,** autocorrelation analyses in roots before and after grafting.

**Supplementary Fig. 4.**
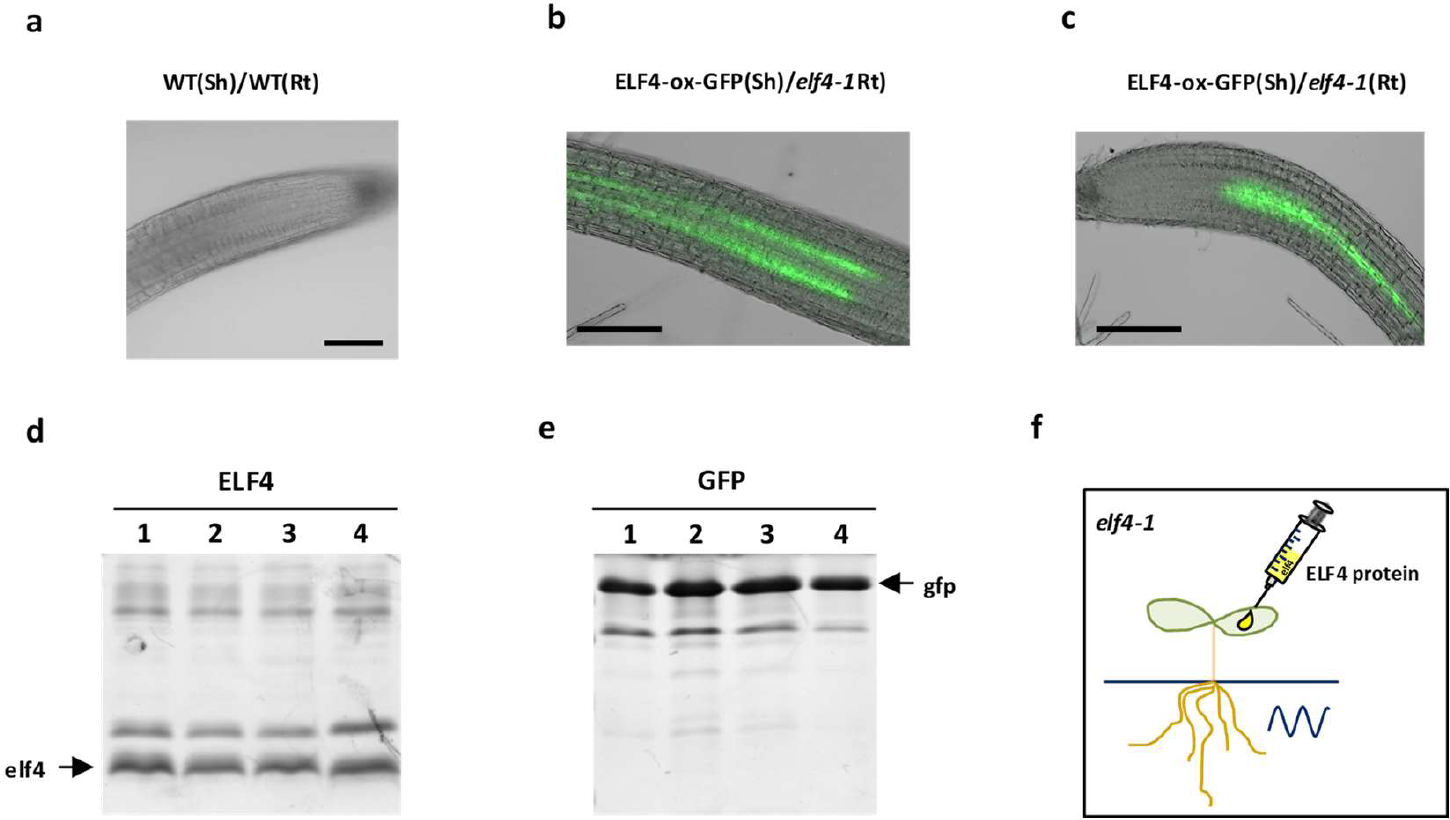
Analyses of ELF4 movement from shoots to roots. **a,** Representative image showing the lack of fluorescence signals in roots of WT scion and WT rootstock. **b, c,** Representative images showing fluorescence signals in roots of ELF4-GFP-ox scion and WT rootstock. Scale bars: 100 μm. **d,** ELF4 and **e,** GFP proteins purified from bacteria and **f,** injected in shoots of *elf4-1* mutant plants to examine rhythmic recovery in roots. At least two biological replicates were performed per experiment.

**Supplementary Fig. 5.**
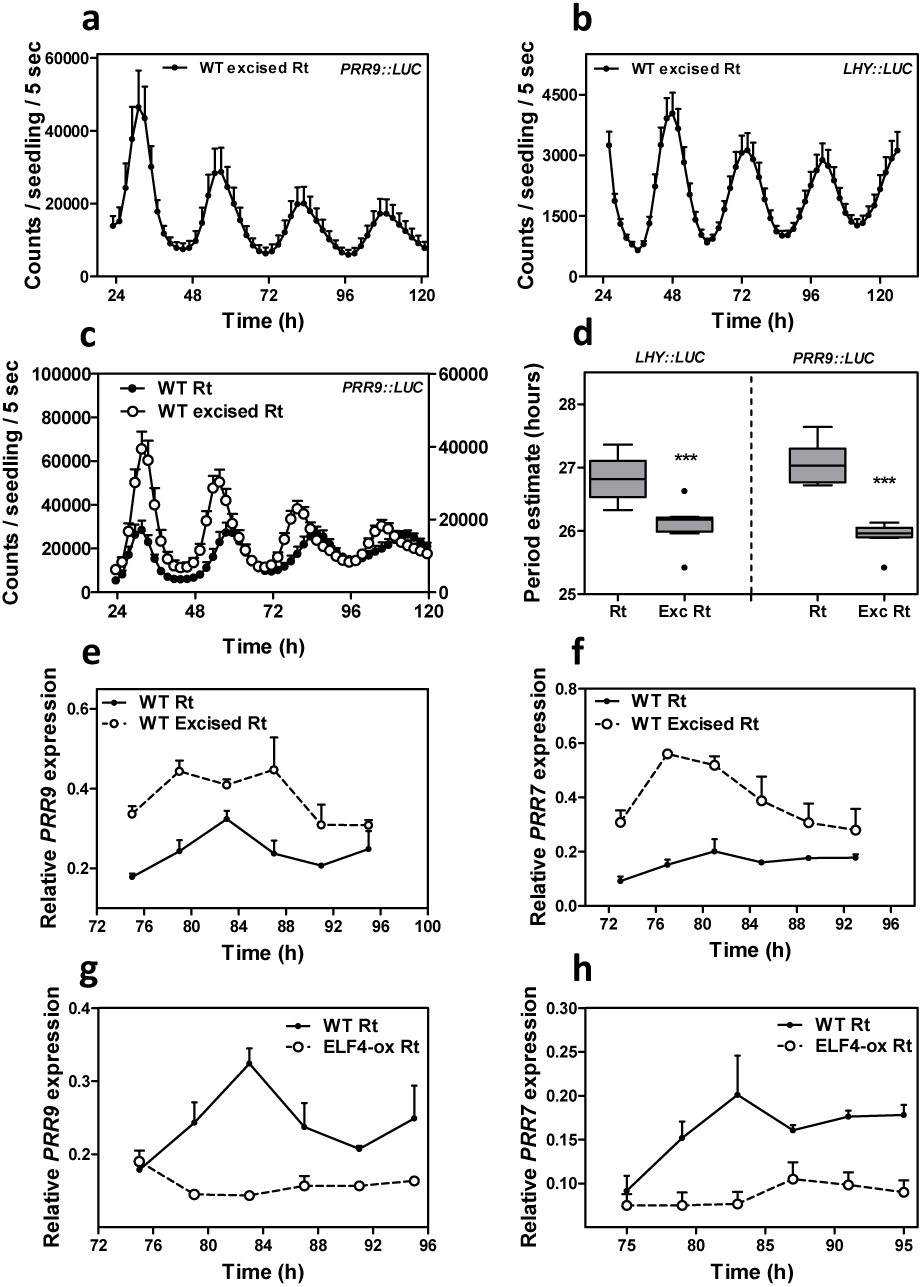
Shoot excision advances the phase and shortens the circadian period in roots. Luminescence of **a,** *PRR9∷LUC* and **b,** *LHY∷LUC* circadian rhythms in WT excised roots. **c,** Comparison of *PRR9∷LUC* circadian rhythms in WT intact versus excised roots. **d,** Period estimates of *LHY∷LUC* (left graph) and *PRR9∷LUC* (right graph) rhythms in WT intact versus excised roots. *** p-value<0.0001. Circadian time course analyses of **e,** *PRR9* and **f,** *PRR7* mRNA expression in WT intact versus excised roots. Circadian time course analyses of **g,** *PRR9* and **h,** *PRR7* mRNA expression in WT and ELF4-ox intact roots. The mRNA expression and promoter activity analyses were performed under constant light conditions previous synchronization of plants under LD cycles at 22°C. At least two biological replicates were performed per experiment.

**Supplementary Fig. 6.**
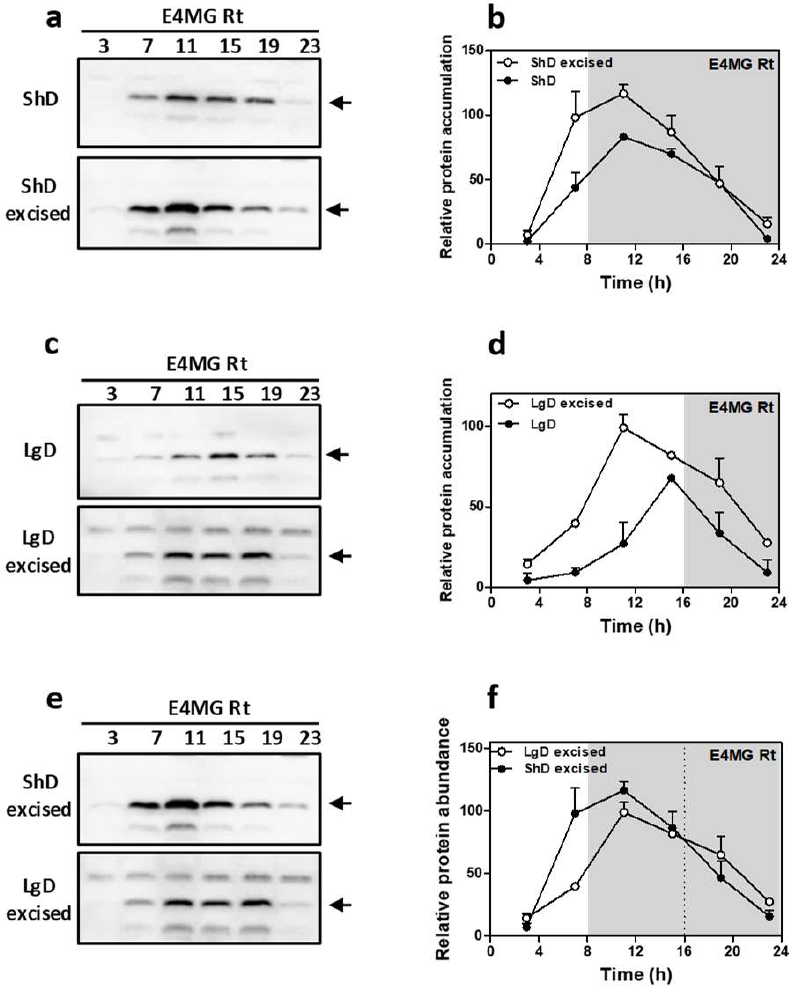
Excision advances the phase of ELF4 protein accumulation in roots under entraining conditions. **a,** Western-blot analysis and **b,** quantification of ELF4 protein accumulation in ELF4 Minigen (E4MG) intact and excised roots under ShD (also in Fig. 3c-d). **c,** Western-blot analysis and **d,** quantification of ELF4 protein accumulation in E4MG intact and excised roots under LgD (also in Supplementary Fig. 3e-f). **e,** Western-blot analysis and **f,** quantification of ELF4 protein accumulation in E4MG excised roots under ShD and LgD (also in Supplementary Fig. 3e-f). At least two biological replicates were performed per experiment.

**Supplementary Fig. 7.**
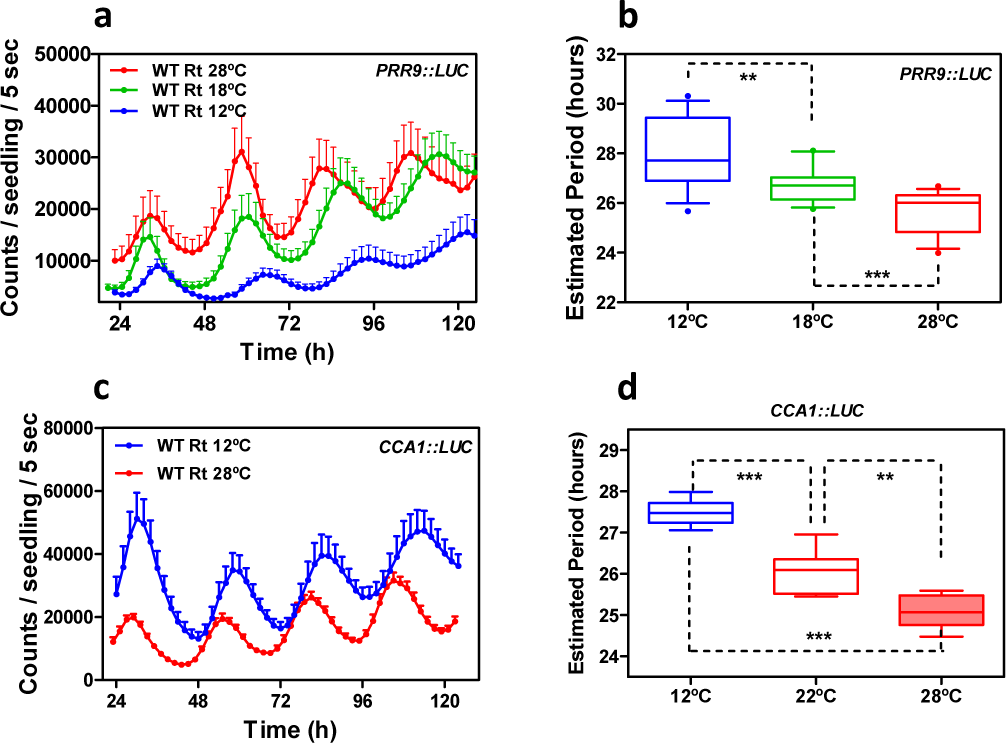
The root clock is not temperature-compensated. **a,** Luminescence of *PRR9∷LUC* rhythmic oscillation in roots at 28°C, 18°C and 12°C. **b,** Circadian period estimates of *PRR9∷LUC* rhythms at 28°C, 18°C and 12°C. **c,** Luminescence *CCA1∷LUC* rhythmic oscillation in roots at 28°C and 12°C. **d,** Circadian period estimates of *CCA1∷LUC* rhythms at 28°C, 22°C and 12°C. *** p-value<0.0001; ** p-value<0.005. At least two biological replicates were performed per experiment.

**Supplementary Fig. 8.**
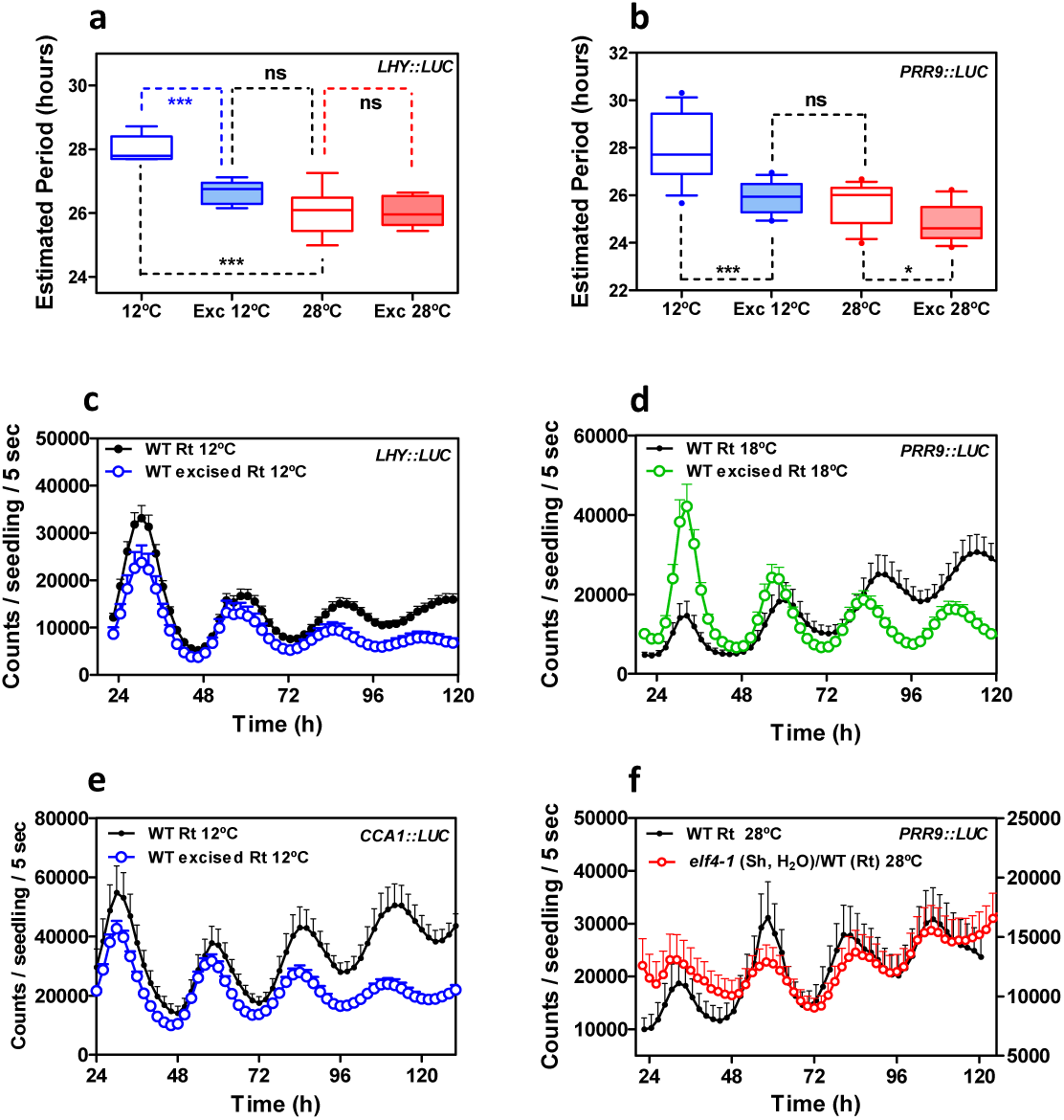
Circadian rhythms in excised roots at various temperatures. Circadian period estimates of **a,** *LHY∷LUC* and **b,** *PRR9∷LUC* in intact versus excised WT roots at 12°C and 28°C. *** p-value<0.0001; * p-value<0.05; ns: non-significant. **c,** Luminescence of *LHY∷LUC* rhythmic oscillation in WT intact and excised roots at 12°C. **d,** Luminescence of *PRR9∷LUC* rhythmic oscillation in WT intact and excised roots at 18°C. **e,** Luminescence of *CCA1∷LUC* rhythmic oscillation in WT intact and excised roots at 12°C. **f,** Luminescence of *PRR9∷LUC* rhythmic oscillation in roots of WT and *elf4-1* scion into WT roots at 28°C. Water instead of luciferin was added to the wells containing *elf4-1* scions. Promoter activity analyses were performed under constant light conditions previous synchronization of plants under LD cycles at 22°C. At least two biological replicates were performed per experiment.

